# Phylogenomic Reconstruction Supports Supercontinent Origins for *Leishmania*

**DOI:** 10.1101/028969

**Authors:** Kelly M. Harkins, Rachel S. Schwartz, Reed Cartwright, Anne C. Stone

## Abstract

*Leishmania*, a genus of parasites transmitted to human hosts and mammalian/reptilian reservoirs by an insect vector, is the causative agent of the human disease complex leishmaniasis. The evolutionary relationships within the genus *Leishmania* and its origins are the source of ongoing debate, reflected in conflicting phylogenetic and biogeographic reconstructions. This study employs a recently described bioinformatics method, SISRS, to identify over 200,000 informative sites across the genome from newly sequenced and publicly available *Leishmania* data. This dataset is used to reconstruct the evolutionary relationships of this genus. Additionally, we constructed a large multi-gene dataset; we used this dataset to reconstruct the phylogeny and estimate divergence dates for species. We conclude that the genus *Leishmania* evolved at least 90-100 million years ago. Our results support the hypothesis that *Leishmania* clades separated prior to, and during, the breakup of Gondwana. Additionally, we confirm that reptile-infecting *Leishmania* are derived from mammalian forms, and that the species that infect porcupines and sloths form a clade long separated from other species. We also firmly place the guinea-pig infecting species, *L. enrietti*, the globally dispersed *L. siamensis*, and the newly identified Australia species from kangaroos as sibling species whose distribution arises from the ancient connection between Australia, Antarctica, and South America.

## Introduction

*Leishmania* is a genus of parasitic trypanosomatid protozoa responsible for the human disease complex, leishmaniasis, which is estimated to cause the ninth largest disease burden among infectious diseases. *Leishmania spp.* are transmitted primarily by sandflies to human hosts and animal reservoirs. Leishmaniasis is endemic to poverty-stricken countries, is rising in incidence by nearly two million cases annually, and lacks effective treatment or vaccine ^1^. At least twenty *Leishmania* species transmitted via insect vector cause disease in humans with three main clinical manifestations: visceral, cutaneous and mucocutaneous. Despite extensive work since its first description in 1903, conflicting hypotheses persist regarding the evolution of *Leishmania* ^2-6^. This lack of consensus regarding current species relationships is due to incongruence between molecular and non-molecular data, as well as among molecular studies ^7,8^. Additionally, available sequence data are skewed towards human-infecting species; comparatively scarce data from wild reservoir/host populations limits evolutionary reconstructions.

Thus, the majority of molecular phylogenetic studies of *Leishmania* have relied on a few genetic loci to infer evolutionary relationships ^15^. These limited data may not accurately reflect the true history of the genus and also provide limited information with which to estimate the age of splits between clades ^9,16^. The evolutionary history of this genus can be resolved with larger molecular datasets and by using these data to estimate the timing of the splits among clades within the phylogeny ^9-14^.

### Proposed origins hypotheses

*Leishmania* are parasites that require two species for a complete life cycle: an insect vector and vertebrate host. Before the development of molecular biology techniques, the genus was divided into two subgenera based upon where the parasite developed within the vector (midgut vs. hindgut) ^17^. These subgenera are *Leishmania*, consisting of all Old World species and one species complex found in the Americas, and subgenus *Viannia*, comprised exclusively of New World species. The species that infect Old World reptiles have since been placed in the subgenus *Sauroleishmania* ^18^ while many other *Leishmania* species, including newly described lineages, remain unclassified.

The biogeography of vectors and distinct vertebrate hosts informs hypotheses for the origins of the genus. Three hypotheses have been proposed for the origin of *Leishmania*. The Palaearctic hypothesis assumes an origin in Cretaceous lizards with recent migrations to the Nearctic and Neotropics across land bridges. This hypothesis suggests that *Sauroleishmania* form a clade that is sister to all other species and is supported by non-molecular data, namely host-phylogenies, biogeography, and evidence of an ancestral parasite, *Paleoleishmania*, found fossilized in Cretaceous amber in modern-day Burma that is similar to species restricted to New World mammals^7,8,19-21^.

Alternatively, *Leishmania* has been hypothesized to originate in the Neotropics. This hypothesis is supported by sequence-based phylogenies ^4,10-12,22^. In these phylogenies, New World species are ancestral to those found in the Old World. Thus, the reptile-infecting forms are derived from mammalian forms of the parasite rather than being ancestral to them, contrary to the Palearctic hypothesis. The Neotropical hypothesis, however, requires two separate intercontinental migrations of the parasites: first the ancestor of *Leishmania / Sauroleishmania* to the Old World, followed by the migration of a member of the *Leishmania* subgenus back to the New World. Alternatively, there might have been two migrations into the Old World: once by an ancestor of *Sauroleishmania* and second by the ancestor of the Old World *Leishmania* species. The timing of these migrations must have coincided with appropriate land bridges.

Finally, the multiple origins hypothesis suggests division of *Leishmania* into two lineages on Gondwana ^9^. One lineage led to the subgenera *Leishmania, Viannia,* and *Sauroleishmania* (termed Euleishmania), while the second diverged into all other species, which to date are restricted to New World mammals (Paraleishmania) ^4^. The breakup of Gondwana subsequently separated the ancestors of the *Viannia* subgenus and the *Leishmania* and *Sauroleishmania* subgenera in South America and Africa, respectively. The amount of diversity observed between the *Leishmania* and *Viannia* subgenera has been cited as evidence for vicariance due to the separation of Africa from the Neotropics ^23^. *L. Leishmania spp.* (e.g. *L. Leishmania amazonensis*) were subsequently brought from Africa through Eurasia to North America. Due to the insect vector’s short lifecycle and weak flying ability this process must have occurred when conditions were conducive to vector survival, and the migration of vertebrate hosts ^24^. The most recent date of an introduction of *Leishmania* subgenus into the Neartic via Beringia would have been the mid-Miocene when temperatures were warm enough for sandfly survival ^13^.

This study represents the first phylogenomic analysis of *Leishmania*, employing over 200,000 variable sites and 49 genes from across the genome. This large dataset counters previous challenges suggesting that different substitution rates in different genes can confound estimates of relationships among *Leishmania* ^9^. The size of the dataset also allows us to estimate the timing of divergence among clades, which until now has been purely speculative. Our results support the hypothesis that *Leishmania* clades separated prior to, and during, the breakup of Gondwana. Additionally, we confirm that reptile-infecting *Leishmania* are derived from mammalian forms, and that the species that infect porcupines and sloths form a clade long separated from other species. We also firmly place the guinea-pig infecting species, *L. enrietti*, the globally dispersed *L. siamensis*, and the newly identified Australia species from kangaroos as sibling species whose distribution arises from the ancient connection between Australia, Antarctica, and South America.

## Materials and Methods

### Whole genome shotgun sequencing

We sequenced whole genomes of twelve species of *Leishmania* and one species of *Endotrypanum* (Table 1). Organisms were grown as promastigotes ^25^ at 22°C in Schneider’s Medium supplemented with 20% heat inactivated FCS and 17.5 mg/mL gentamycin. The cells were pelleted and washed twice in phosphate-buffered saline (PBS) and gDNA extracted according to established methods. The cells were incubated in lysis buffer (100mM NaCL, 10mM TrisCl pH 8.0, 0.5mM EDTA, 0.5% SDS, 0.1mg/ml fresh Proteinase K) for 12-18hrs at 50C, and purified via phenol/chloroform/isopropyl alcohol. Extract was incubated in RNase A (10mg/ml) for 2.5hrs at 37C and dialyzed overnight at 4C with 3 changes of PBS buffer. DNA concentration was evaluated with spectrophotometry (Beckman Coulter DU730) and re-extracted if contaminated with phenol. Paired-end reads (100 bp) were sequenced on an Illumina HiSeq2000 by the University of Arizona Genome Core. The number of pairs of reads for each species ranged from 2.1-14.3 million, or 10 - 67x average coverage for the ~34 million bp genome.

Additional shotgun genomic data were downloaded from the European Nucleotide Archive (ENA; Table S1). These data represent all publicly available genome data per *Leishmania* taxon.

### Phylogenetically informative data

#### Variable sites from across the genome

Although reference genomes are available for *Leishmania,* initial attempts to align shotgun sequencing reads to these references resulted in low alignment rates. Therefore, to extract phylogenetically informative data from the available shotgun sequences, we used the bioinformatics pipeline SISRS ^26^. This method identifies sites that are fixed for each species and variable across species to construct a multiple-species alignment. SISRS determines the nucleotide each species has at a site via strict consensus. If this fails or there is no data for that site in a species, the information is considered “missing”. We produced dataset VS-m6 allowing up to six taxa to have missing information per site (missing data were allowed to ensure sufficient data to determine the phylogeny). To ensure that linked sites would not bias our results, we subsampled this dataset by sampling only one site per sequence fragment, producing dataset VS-m6s. Finally, we produced an additional alignment, VS-t80, with no missing data allowed, but with a lower calling threshold of 0.8, i.e. 80% of bases for that taxon must be one allele.

#### 2.2.2 Additional gene data

Because branch length estimates can be incorrect for variable site datasets, we developed an additional dataset of 49 genes with putative or known function. This dataset also allowed us to compare the results of two approaches to estimate the *Leishmania* phylogeny. We first obtained these genes from the *L.V. braziliensis* MHOM/BR/75/M2904 or M2903 reference genome (Table 2). The sequence of each gene for all other species was then extracted from the shotgun sequencing reads using the following pipeline: (1) Reads from each species were aligned to the reference genes using Bowtie2 ^27^. (2) We used the mpileup feature of SAMTools^28^ to obtain the information from each read for each site. (3) The base for each site was identified based on whether at least 80% of the reads at that site contained a single base. (4) Alignments were adjusted using MAFFT ^29^ with default settings. This pipeline is now automated as a part of SISRS.

We then added to the gene dataset all data available for two recently described *Leishmania sp.:* an isolate from Australian kangaroos (strain AM-2004) ^30^ and *L. siamensis ^31,32^*. These data were downloaded from GenBank. For AM-2004 we obtained partial sequences of three loci. For *L. siamensis* we obtained partial sequences of six loci, which includes two genes (Table S2).

### Phylogenetic analysis

#### Concatenated variable site analysis (3 datasets)

The practice of concatenating thousands of likely unlinked sites into a single alignment ^33^ can elide the variable genealogical histories of different chromosomal regions. However, variable sites cannot be partitioned by linkage to separate the history of genes from the history of the species, and it is computationally challenging to consider each site separately for a dataset of this size. We thus treated the SISRS variable site data as a single concatenated locus ^34^. For each dataset, we constructed a phylogeny using maximum likelihood (ML) in RAxML 8.0.20 ^35^ with a General Time Reversible (GTR) model and substitution rates following a discrete gamma distribution with four categories for 1000 bootstrap replicates and allowing for ascertainment bias correction using the Lewis model ^36^.

#### Gene data

For coding genes, partitioning by codon position was found to provide a better fit than by gene alone ^37^. The best way to partition the data was determined by the program *partitionfinder* with default settings ^38^. Input partitions were separate codon positions for each gene. We employed the Bayesian Information Criterion for model selection to avoid overparameterization. The resulting 20 partitions were used to estimate the phylogeny in a ML framework; this analysis was implemented in RAxML 8.0.20 ^35^ using a GTR model with gamma distributed rate heterogeneity across sites, for 100 bootstrap replicates. This analysis was repeated five times with random starting seeds to identify the ML tree and avoid local optima.

#### Individual gene analysis

ML trees for each gene were constructed identically in RAxML with 1000 bootstrap replicates. Although concatenated multi-gene datasets are found to be robust in phylogenetic inference, even without explicitly accounting for large variations in rates, lengths and GC content, systematic biases in the dataset can lead to high support of the wrong tree ^39^. We follow Gadagkar, et al. ^39^ recommendation to report the gene support frequency for each given partition. A majority rule (MR) consensus tree was generated in RAxML from the best scoring ML trees of individual gene trees for which we had all ingroup taxa. When necessary the outgroup *Crithidia* was pruned using Newick Utilites ^40^.

#### Estimating divergence time

We used two approaches to estimate the timing of divergences among clades. Time estimates allow us to evaluate the plausibility of the proposed hypotheses for the origin of *Leishmania*. Due to poor fossil preservation of *Leishmania,* no secure fossil calibration dates within the genus exist. Thus, our first approach was to add two outgroup species, *Trypanosoma cruzi* and *T. brucei*, to the concatenated gene dataset. The divergence date for these species is believed to be 100 million years (my) ago based on the timing of the split between Africa and South America ^14,41^. Using our known tree for *Leishmania* with these two additional species, divergence times were estimated using RelTime ^42^.

Our second approach was to use a calibration date of 40 million years for the split between *L. enriettii, L. siamensis* and *Leishmania sp. Ghana* with the Australian isolate *Leishmania* sp. AM-2004, which corresponds to the breaking of the connection between South America and Australia via Antarctica. In this analysis we only used the sequences available for AM-2004 to avoid any effect of missing data on branch lengths. For this reason, we were able to include additional *Leishmania* taxa for which these few loci were also publicly available (Table S3). We constructed dated phylogenies using BEAST v1.8.2 ^43^ with two unlinked partitions, one for 18s/ITS/5.8s and the other for RNA polymerase II large subunit. We estimated substitution models in jmodeltest^44^, and given those results, implemented a TN93 substitution model with rates estimated from a gamma distribution with four rate categories and a relaxed lognormal molecular clock for 18,000,000 generations (high ESS values and convergence were observed in Tracer 1.5 ^45^ at this point). A normally distributed prior with a mean of 40 my was specified for the node leading to Australia kangaroo isolate, Leishmania sp. AM-2004.

## Results

### Phylogenetic analyses

#### Variable sites data

The number of variable sites identified for each dataset were 215,644 (VS-m6), 18,312 (VS-m6s), and 2,790 (VS-t80). Based on an alignment with the reference genome for *L. tarentolae*, we determined that these sites are distributed across the *Leishmania* genome. GC content was over 70%, which is higher than the GC content estimated previously of 50–60% (Peacock et al. 2007). The ML phylogeny for all of these datasets supports the *Leishmania* and *Viannia* subgenera as monophyletic clades; *Sauroleishmania* is also monophyletic, although only two taxa were available for phylogenetic analysis, and sister to the clade comprising the other two subgenera (Figure 1). *L. enrietti* is supported as sister to all other Euleishmania. *L. hertigi*, *L. deanei*, and *Endotrypanum* (Paraleishmania), form a clade that is sister to all other *Leishmania* lineages (Euleishmania).

**Figure 1:**
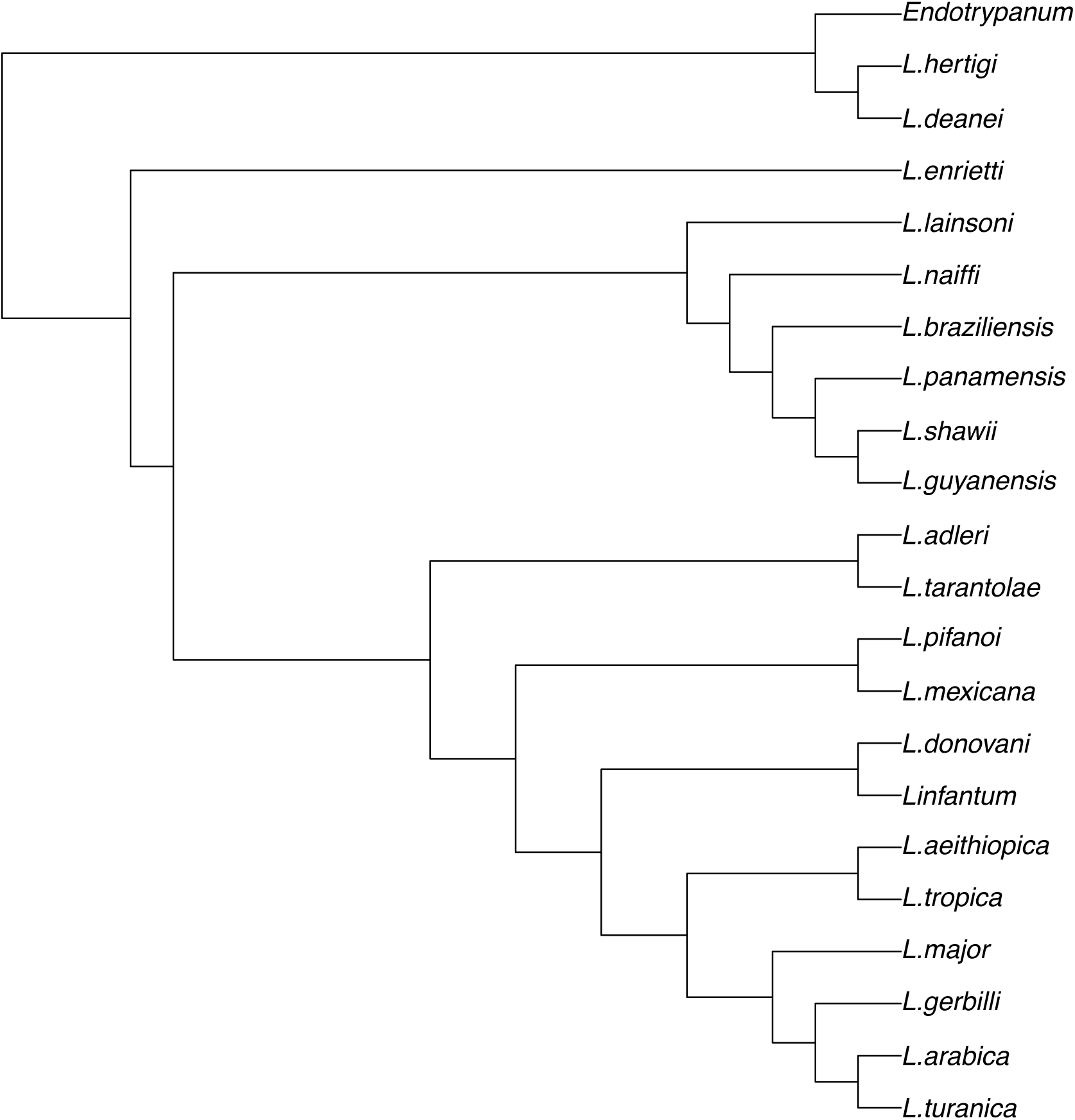
Maximum likelihood tree based on 215,644 variable sites from across the genome. Data were identified using SISRS ^26^. Up to six samples were allowed to have data missing or be ambiguous for any given site. The phylogeny was constructed with RAxML 8.0.20 ^35^ using the correction when only variable sites are present ^36^. Bootstrap support values are 100 at every node. Identical results were obtained using a dataset containing only one site per sequence fragment, and one with no missing data allowed, but with a lower calling threshold of 80%.

#### Partitioned genes

A total of 70,447 sites in 49 genes were concatenated in the alignment. The resulting ML tree is identical to that for the variable sites data, with the exception of the placement of *L. enrietti* as sister to the clade containing the subgenera *Leishmania* and *Sauroleishmania*, rather than as sister to Euleishmania (Figure 2). However, this placement was supported by less than 50% of individual gene trees. Interestingly, when ML trees were constructed separately from each codon position the phylogeny was identical to that constructed for the variable site data, although the third codon position phylogeny had low support at several nodes. *Leishmania* AM-2004 and *L. siamensis* are most closely related to *L. enrietti*.

**Figure 2:**
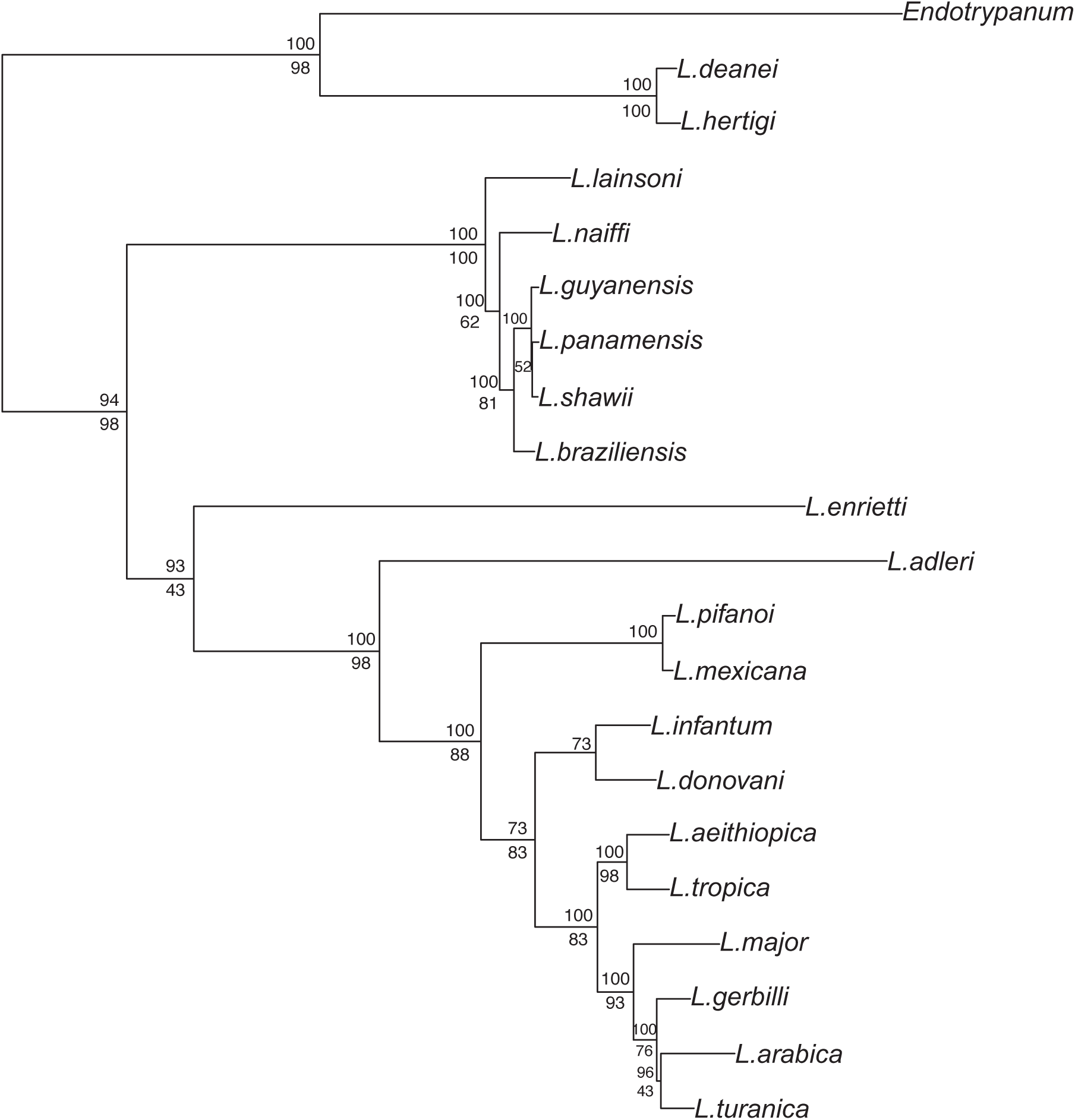
Maximum likelihood tree of *Leishmania* based on 49 genes. The phylogeny was constructed with RAxML 8.0.20^35^; data were partitioned by gene and codon position, with individual partitions grouped together using partitionfinder^38^. Top and bottom node labels indicate bootstrap support from 100 replicates and the percent of genes that support that node when the phylogeny for each gene was calculated individually, respectively.

### Divergence time estimates

Divergence dates were first estimated using the gene dataset for *Leishmania* and two species of *Trypanosoma,* and Reltime with a 100 my calibration point at the split between two outgroup species, *T. cruzi* and *T. brucei* (Figure 3). The estimated date for the split of *L. donovani-L. major* was 24.2 mya,

**Figure 3:**
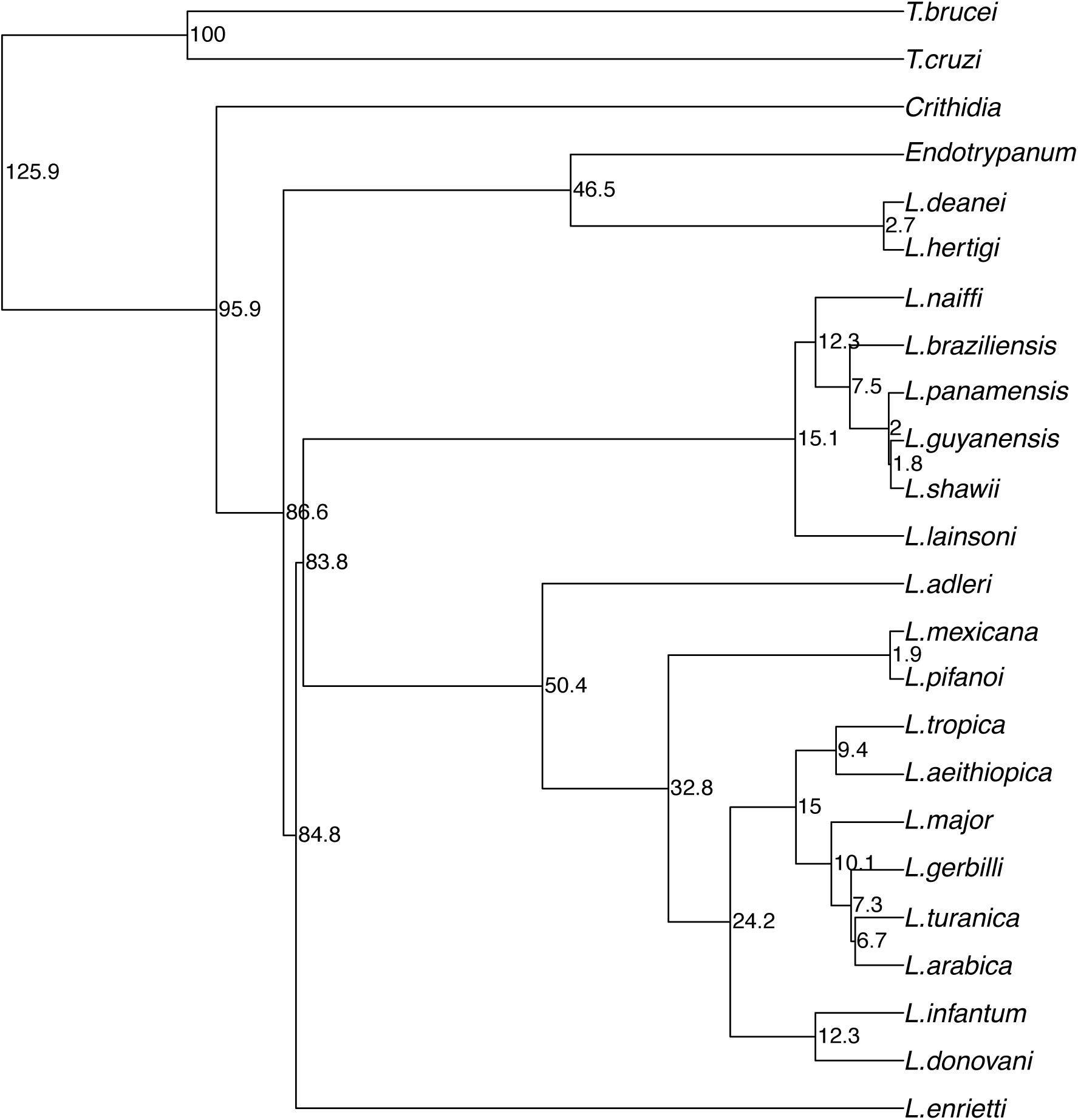
Dated phylogeny based on 49 genes. Constructed with Reltime using the calibration date of 100 mya for the split of *Trypanosoma brucei* and *T. cruzi.* Values at nodes are in million of years.

We were unable to obtain genome data for the sister genus to *Leishmania*, which is *Leptomanas*; thus, we approximate the origin of *Leishmania* based on the timing of the split of the genus into two clades, and the divergence of *Leishmania* and *Crithidia.* These dates lead to an approximate origin of 90mya.

Secondly, divergence dates were estimated using available loci for the Australian species in BEAST v1.8.2^43^ and setting a normal prior distribution of 40 my on the node ancestral to this isolate and other members of the “*L. enriettii* complex”. In this case we estimated the median date of the split of *L. donovani-L. major* to be 21.2 (15.2–47.3 95% HPD) mya. These results suggest an older origin of the genus between 140 and 119 mya (i.e. between the split between *Leptomonas* and *Leishmania*, and the split between Paraleishmania and Euleishmania) (Figure 4 and Figure S1).

**Figure 4:**
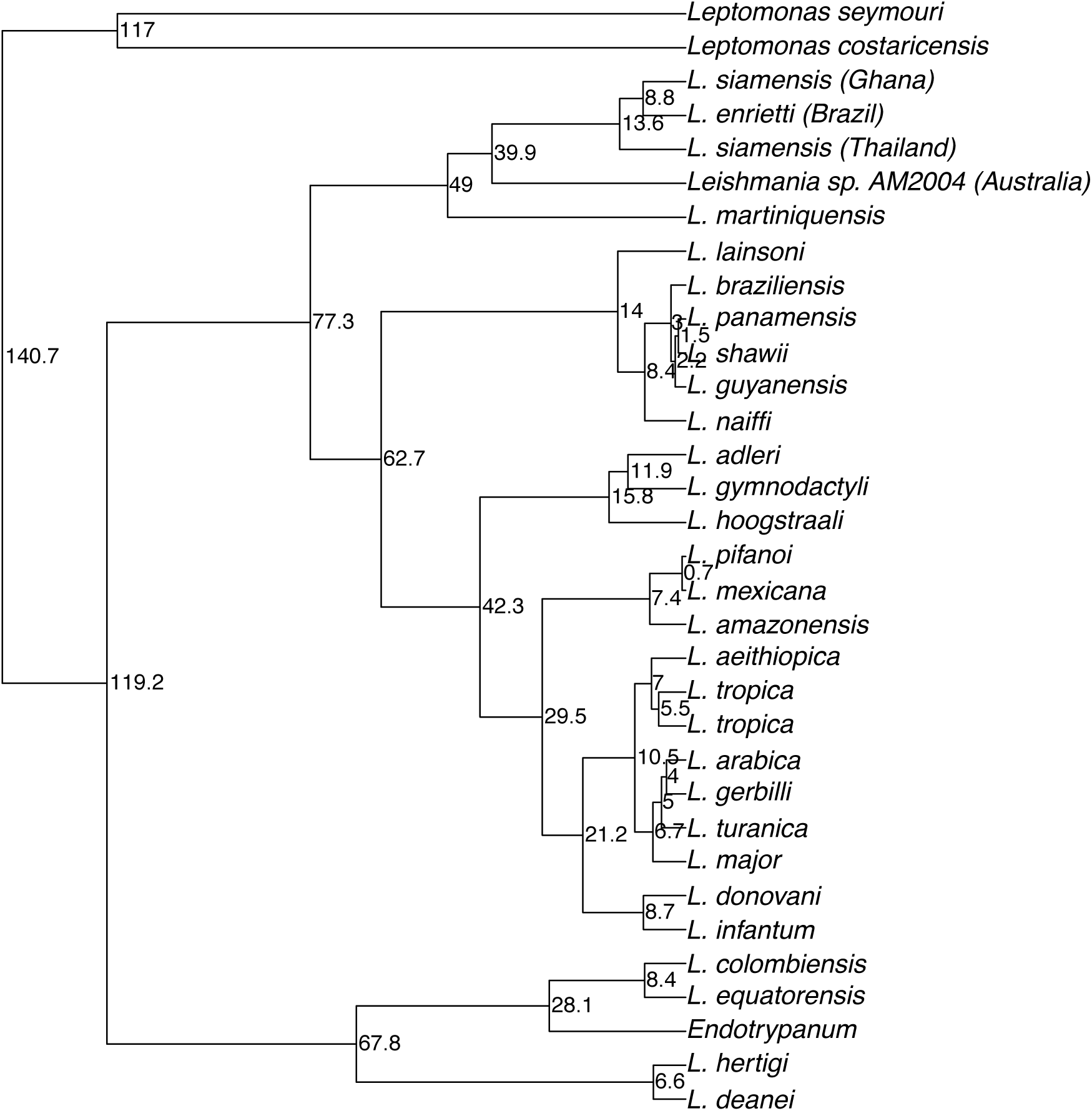
Phylogeny constructed using the loci available for *L. sp.* AM-2004 in BEAST with a relaxed molecular clock. Node values denote millions of years before present. See Figure S1 for posterior probability and 95% HPD error bars. This phylogeny places the Australia sample as sister to *L.siamensis* and *L. enrietti*. Setting the split between the Australia sample and those taxa at 40 mya (a minimum time for the loss of connection between Australia and South America via Antarctica) allows us to estimate dates for the rest of the phylogeny. In particular, we estimate the median age of the split between *Leishmania* and *Leptomonas*, its sister genus, to be at 119 mya, and the split between the subgenera *Sauroleishmania / Leishmania* and *Viannia* at 62.7 mya. This result suggests that the *Leishmania* genus emerged prior to the breakup of Gondwana, and the latter split was caused by the separation of Africa from South America.

## Discussion

### Data

Shotgun sequencing data of thirteen *Leishmania* isolates (this study) and eleven species available publicly enabled us to mine whole genome datasets for hundreds of thousands of phylogenetically informative sites. Previous molecular datasets used for phylogenetic reconstruction ranged from 1–31 genes ^47^. Rather than relying on a few known genes, these data were sampled across the genome. The SISRS method is particularly valuable for *Leishmania sp.*, which like many other non-model organisms, do not have a suitable reference genome for this purpose.

### Phylogeny

Our results place species of Old and New World subgenus *Leishmania* in a monophyletic clade separate from species of the exclusively New World subgenus *Viannia*. These results are consistent with other molecular-based trees, including those that place *Sauroleishmania* as sister to the *Leishmania* subgenus ^12,48^.

Our genome-wide, variable-site tree places the *L. enrietti* genome, a parasite of the New World guinea pig, sister to all Euleishmania. This result is consistent with some prior molecular phylogenies ^2,10,22^; however, it differs from early hypotheses that suggest *L. enriettii* is a member of the New World *L. Leishmania* subgenus ^49,50^ or later work suggesting it is sister to the entire genus^20,51^.

When all codon positions are considered, the concatenated gene dataset produces a conflicting topology whereby *L. enrietti* is not basal to all Euleishmania but only to the *Sauroleishmania / Leishmania* subgenera. This conflict is not necessarily surprising given the short branches leading to the splits of *L. enrietti* and the *Euleishmania* clades, which often reflects incomplete lineage sorting and would lead to different relationships estimated from different genes. Adding more loci, as we have done with the variable site dataset, is advocated in this case to increase resolution ^52^. It is interesting to note that strong support for the tree produced from the multi-gene data is obtained when using only the first and second codon position of the gene data, and lower support for this same topology results from separate analysis of third positions. Substitutions are expected to be most rapid at the third position ^53^. At these time scales, saturation in the third position may negatively affect the phylogenetic signal, as reflected in the decrease of support.

As with nearly all other analyses to date, the molecular evidence does not support the taxonomic classification of *Endotrypanum* as a separate genus. Relative to molecular markers, there are few morphological characters used to classify unicellular organisms, calling their utility into question, when compared to molecular data ^54^. Sequence data support their inclusion in the *Leishmania* genus, unless all other Paraleishmania species are misclassified.

### Dates

The calibration of 40 mya for the split between the “*L. enriettii* complex” and the Australian isolate *Leishmania* sp. AM-2004, like other fossil dates, provides only a minimum time since two species diverged. However, an examination of the resulting dates on other nodes within the tree, namely the split to *L. donovani* and *L. major*, is consistent with previously published dates. The dates for the split of *L. donovani* and *L. major* calculated using both datasets (24.2 and 21.1 mya respectively), with different calibration points, compare favorably to those published previously (24.6–14.7 my) ^14^. This result suggests that results at other nodes may be considered reasonably valid when evaluating hypotheses for the origin of *Leishmania*.

### Origins hypotheses

#### The Paleoarctic hypothesis

Our results reject the Palaearctic origin hypothesis for *Leishmania^7,20^*. First, *Sauroleishmania* are not sister to all other *Leishmania,* as would be suggested by this hypothesis. Second, our estimated dates conflict with a Pliocene introduction of the subgenus *Viannia* to the Neotropics at ~5 mya after the reformation of the Panama isthmus. If Kerr ^7^ is correct in asserting that reptiles were the first hosts of *Leishmania* in the Cretaceous, extinction events associated with the K-T boundary could have erased evidence of those lineages; however, this speculation does not affect our interpretation.

#### The Neotropical hypothesis

Our results also allow us to reject the Neotropical Origins hypothesis. Under this hypothesis, the current global distribution of *Leishmania* is a purely a result of dispersal through vector / reservoir migration and not vicariance. Lukes et al.’s (2007) version of the hypothesis proposes the ancestor of Paraleishmania evolved in the Neotropics between 46–36 mya and dispersed north. Similarly, Noyes et al. (1997) and Noyes (1998) propose that the Leishmania / Endotrypanum clade evolved in the Neotropics between 65–40 mya and dispersed to the Nearctic and to the Palearctic through land bridges; the Neotropics however were isolated at this time. Our results suggest an origin of the clade at least 90 mya, which is not consistent with these proposed dates.

#### The Multiple Origins hypothesis

The Multiple Origins scenario refers to the separation and subsequent independent evolution of the *Viannia* and *Leishmania* subgenera before 90-100 mya. This is the only formalized hypothesis that includes vicariance as a mechanism for the evolution of the major *Leishmania* clades. However, the authors are unclear about the emergence of Paraleishmania; while this group is clearly shown as diverging from Euleishmania prior to the separation of *Viannia* and *Leishmania* subgenera, they state that Paraleishmania migrated to the New World with the introduction of hystricomorph rodents (e.g. porcupines) in the “early Cenozoic”. This explanation is unlikely because hystricomorph rodents do not appear in the fossil record until 23 mya (Paleobiology database, fossilworks.org).

We propose a modification and expansion of the multiple origins hypothesis, which we now term the Supercontinent hypothesis, for the origin of the *Leishmania* genus. In this scenario, the ancestor of *Leishmania* emerged from monoxeneous parasites (those found in a single host) on Gondwana ^51^. As with the Multiple Origins hypothesis, the ancestors of Paraleishmania, and the *Viannia* and *Leishmania* subgenera emerged before separation of Gondwana ~90-100 mya and possibly already sustained a global distribution, as suggested by the placement and divergence of strains found in Asia and Africa, here *L. siamensis*, *L. martiniquensis*, and *Leishmania* sp. Ghana. The Supercontinent proposal is in agreement with speculations proposed by Shaw^55^ who suggested that an adaptation to mammals occurred around 90 mya when mammals began to radiate and Africa became fully isolated. Additionally, levels of genetic diversity between the *Viannia* subgenera and *Sauroleishmania / Leishmania* subgenera have been cited previously as a reflection of vicariance after the separation of South America and Africa^23^. These speculations are consistent with the estimated divergence dates presented here using two methods with two types of datasets, one genome-wide and one gene-based, with separate calibration information, respectively.

Only one migration of a lineage in the *Leishmania* subgenus back to the New World is required by this hypothesis (notwithstanding the historical transfer of *L. infantum* to the New World by European settlers, often termed *L. infantum chagasi ^56,57^*). The global distribution of the phlebotomine sandfly genera that almost exclusively serve as vectors for the parasite likely resulted from the breakup of Pangaea and subsequent continental splintering ^58,59^.

Our results are consistent with Early Cretaceous fossils of *Paleoleismania proterus* found in sandflies trapped within Burmese amber ~100 mya, which are reportedly evidence of the first digenetic trypanosomatids ^21^. There is no way to confirm the genus but the organisms are morphologically similar to *Leptomonas*, the sister of *Leishmania*. Interestingly, this species is believed to be associated with reptile hosts.

The recent discovery of *L. siamensis* and other linages that fall within the “*L. enriettii* complex”, a clade basal to all Euleishmania, in humans and other mammals in North America, Europe, West Africa and Asia further highlights the plausibility of an ancient global dispersal predating continental splintering; because few data are currently available for all lineages within the new *L. enriettii* clade, further work is necessary to rule out recent introduction of *Leishmania* to those regions. However, the placement of AM-2004 as sister to lineages found in the New World (*L. enriettii*) is consistent with the biogeography of other groups separated by the breakup of Australia, Antarctica and southern South America (e.g. the plant genus *Nothofagus)*. The Supercontinent hypothesis for *Leishmania* genus also draws parallels with a related kinetoplastid parasite, *Trypanosoma*. Hamilton, et al. ^60^ hypothesize a southern-supercontinent origins in which *T. brucei* evolved in Africa and *T. cruzi* in the New World following the breakup of Gondwana. Divergent lineages of *Trypanosoma* has also been found in Australia ^61^, therefore much like the southern-supercontinent hypothesis for *Trypanosoma* evolution, the position of the kangaroo *Leishmania* isolate is critical in our origins scenario. This parallel pattern of evolution between related parasites *Trypanosoma* and *Leishmania* potentially suggests similar evolutionary processes.

The most ancestral lineages of *Leishmania* — those crucial to resolve the evolutionary history of the genus — are not only those typically found in wild reservoir populations, but also the samples for which fewest data exist. Animal reservoirs are critical for maintaining *Leishmania* in the wild. Sampling strategies refocusing on wild host and vector populations, recently urged by many researchers, will shed light on the deepest nodes of the phylogeny and ultimately, on the processes of zoonotic transfer to humans. Until additional NGS data are available from unclassified and newly described taxa, we offer genome wide data and multiple approaches to estimate divergence dates further contributes to our understanding of the evolution of *Leishmania*.

## Acknowledgements

Genomic DNA from *Leishmania* isolates was obtained from the lab of Diane McMahon-Pratt at the Yale School of Public Health and from Lucille Floeter-Winter at the University of Sao Paolo. This work was supported by a National Science Foundation Doctoral Dissertation Improvement Grant [grant number BCS-1232582 to K. Harkins and A. Stone], School of Life Science (ASU) [R. Cartwright], and a National Science Foundation Advances in Bioinformatics Grant [grant number DBI-1356548 to R. Cartwright]. Sudhir Kumar and Jay Taylor provided guidance and feedback on the development of the project.

## Supplementary Material

**Table S1:** Raw genome data downloaded from public online databases.

**Table S2:** Accession information of genes acquired for *Leishmania* sp. AM-2004 and L. *siamensis.*

**Table S3:** Accession numbers for additional taxa included in BEAST dating analysis for RNA polymerase II large subunit (Rpol), 18s, Internal Transcribed Spacer 1 (ITS) and 5.8s. All others taxa derive from WGS data listed in Table 1 and S1.

**Figure S1:** BEAST tree with 95% HPD error bars for estimated divergence dates. Scale in millions of years. Node values are posterior probabilities.

